# Triclosan Exposure is Associated with Rapid Restructuring of the Microbiome in Adult Zebrafish

**DOI:** 10.1101/039669

**Authors:** Christopher A. Gaulke, Carrie L. Barton, Sarah Proffitt, Robert L. Tanguay, Thomas J. Sharpton

## Abstract

Growing evidence indicates that disrupting the microbial community that comprises the intestinal tract, known as the gut microbiome, can contribute to the development or severity of disease. As a result, it is important to discern the agents responsible for microbiome disruption. While animals are frequently exposed to a diverse array of environmental chemicals, little is known about their effects on gut microbiome stability and structure. Here, we demonstrate how zebrafish can be used to glean insight into the effects of environmental chemical exposure on the structure and ecological dynamics of the gut microbiome. Specifically, we exposed forty-five adult zebrafish to triclosan-laden food for four or seven days or a control diet, and analyzed their microbial communities using 16S rRNA amplicon sequencing. Triclosan exposure was associated with rapid shifts in microbiome structure and diversity. We find evidence that several operational taxonomic units (OTUs) associated with the family Enterobacteriaceae appear to be susceptible to triclosan exposure, while OTUs associated with the genus *Pseudomonas* appeared to be more resilient and resistant to exposure. We also found that triclosan exposure is associated with topological alterations to microbial interaction networks and results in an overall increase in the number of negative interactions per microbe in these networks. Together these data indicate that triclosan exposure results in altered composition and ecological dynamics of microbial communities in the gut. Our work demonstrates that because zebrafish afford rapid and inexpensive interrogation of a large number of individuals, it is a useful experimental system for the discovery of the gut microbiome’s interaction with environmental chemicals.

## Introduction

The gut-associated microbiome performs vital functions in the gastrointestinal tract, which prevents colonization with pathogens[1,2], stimulates immune system development and function[3,4], and produces micronutrients utilized by the host[5]. Deviation from normal microbiome structure or function, known as dysbiosis, has been associated with human diseases, including diabetes[6], heart disease[7], arthritis[8], and malnutrition[9]. These findings have inspired investigation into the mechanisms of perturbation of the gut microbiome, which have identified several potent modulators of microbiome composition, including antibiotic therapy[10,11], infection with pathogens[12,13], and diet[14,15]. Establishing a comprehensive catalog of the factors that influence gut microbiome structure and function is an essential first step in designing therapies to treat dysbiosis or prevent its deleterious effects.

Humans are exposed to a diverse array of chemicals on a daily basis through contact with the environment. While some of these exposures may be innocuous or even beneficial (e.g., dietary micronutrients), others have been associated altered host physiology and chronic disease[16,17]. Each chemical also represents a potential source of perturbation for microbial communities that might 1) modulate growth rates or kill microbes, 2) alter nutrient availability, or 3) restructure niche space. Although studies of the impact of environmental chemical exposure on gut microbiome structure and function are limited, the information available implicates their influence over shaping the composition and function of microbial communities. For example, dietary chemicals such as dietary fiber can be fermented by microbes in the gut to produces short-chain fatty acids, which can be anti-inflammatory and influence the permeability of the gut epithelial barrier by enhancing tight junction expression[18]. Conversely, the metabolism of other dietary chemicals, such as L-carnitine, can lead to metabolites that are associated with disease[7]. Heavy metals, such as arsenic, lead and cadmium also perturb microbial community composition and metabolic profiles in mice[19] and are associated with altered immune responses, and gut barrier function[20,21]. Interestingly, many of the environmental chemicals to which humans are exposed are antimicrobial in their action. For example, parabens, sulfites, nitrites, nitrates, and many phenols are preservative agents present in foods, cosmetics, and cleaning supplies. Despite generally being considered safe, some of these antimicrobial compounds have been associated with endocrine disruption[22,23] and inflammatory diseases such as ulcerative colitis[24]. However, the impact of exposure to these compounds, or their derivatives, on the structure and function of the microbiome remains unclear.

Due to the diversity of environmental chemicals that animals are exposed to, experimental systems that enable rapid screening of the effect of their exposure on the microbiome need to be developed. Here, we use zebrafish as a model system to explore the impact of environmental chemical exposure on the animal gut microbiome. Zebrafish were selected as they are a widely used toxicology model [25–27], are easily housed in large numbers, are inexpensive, and afford access to a wide array of high-throughput genomic[28], developmental[29], and physiological experimental tools[30,31]. Additionally, zebrafish are a well established model system of ecological dynamics of gut microbial communities [32–34] and have provided valuable insights into interactions between gut microbes and host immune and metabolic processes[35–37]. These features are useful in microbiome studies as the subtle effects of low concentration toxicant exposure are likely to require large sample sizes to resolve, and the follow-up investigations on the impact of the perturbation on host physiology will be benefitted by access to diverse experimental resources.

We use zebrafish to examine the impact of short-term exposure of an environmental antibiotic, triclosan, on the gut microbial communities of adult zebrafish *(Danio rerio)* using 16S rRNA amplicon sequencing. This polychlorinated phenoxy phenol, which is widely used as an antimicrobial agent in consumer products[38], was selected for a variety of reasons. First, triclosan is readily absorbed through the skin and gastrointestinal tracts, and is excreted in urine, breast milk, and feces[39–41]. Second, exposure to triclosan is associated with endocrine disruption in fish and rats, research implicates its role as a liver tumor promoter[22,42,43], and it can alter inflammatory responses by modulating toll-like receptor signaling[44]. Third, triclosan can disrupt microbial communities. Exposing aquatic- or soil-associated microbial communities to triclosan alters community composition and reduces diversity[45,46]. Moreover, triclosan has been associated with disruption of the gut microbiome in juvenile fathead minnows *(Pimephales promelas)*[47]. Here, we found that triclosan exposure is associated with restructuring of the zebrafish gut microbiome over short time intervals in adult fish. In addition, triclosan exposure was associated with increased correlation between a large number of microbial taxa and broad restructuring of microbial interaction networks. Taken together these results indicate that triclosan exposure disrupts the structure and ecological dynamics of the gastrointestinal tract microbiome of adult zebrafish.

## Materials and Methods

### Animals and triclosan exposure

The use of zebrafish in this research project conducted at the Sinnhuber Aquatic Research Laboratory was approved by the Institutional Animal Care and Use Committee at Oregon State University (permit number: 4696). Adult, male, 5D wild type zebrafish were separated into nine tanks (n=5 fish / tank), and placed in flow-through tanks isolated from the rest of the colony. The nine tanks were randomly assigned to a group (unexposed, four-day treatment, seven-day treatment) and fish were fed a commercial pelletized lab feed (Gemma Micro 300; Skretting, Westbrook, ME USA) containing 100*μ*g/g fish a day triclosan, a dose sufficient to cause endocrine disruption in fish[22], or a control diet that was compositionally identical with the exception that it contained no triclosan. There were three exposure groups, (1) a four-day exposure group, which received the triclosan diet for four days followed by a control diet for 3 days, (2) a seven day group which received triclosan laden food for seven days, and (3) an unexposed control group, which received the control diet for seven days. On the morning of the eighth day the remaining fish in each tank were euthanized by ice water bath immersion. Each fish was then surface sterilized using 70% ethanol and intestinal contents were collected by removing the length of the intestine (esophagus to anus), and then gently squeezing the intestine with forceps to extract contents. The contents were collected in a sterile DNAse free tube and stored at −20°C until processing.

### 16S rRNA amplicon library preparation and sequencing

The MoBio PowerSoil® DNA isolation kit (MOBIO, Carlsbad, CA USA) was used to extract DNA from the intestinal contents samples following the manufacturer’s protocol with the addition of an incubation step of 10m at 65°C immediately before bead beating on the highest setting for 20m using Vortex Genie 2 (Fisher, Hampton, NH USA) and a 24 sample vortex adaptor (MOBIO). Two microliters of purified DNA was then used for input into PCR reaction and the remaining DNA stored at −20°C. Amplification of the 16S rRNA gene was performed as previously described using primers directed against the V4 region[48,49]. Amplicons were visualized using gel electrophoresis to ensure a band corresponding to ∼350 bp was present in each library. Each library was then quantified using the Qubit® HS kit (Life Technologies, Carlsbad, CA USA) according to the manufacturer’s instructions and 200ng of each library were pooled. The pooled library was then cleaned using the UltraClean® PCR clean-up kit (MOBIO) and diluted to a concentration of 10nM. The prepared libraries were then subjected to cluster generation and sequencing on an Illumina MiSeq instrument. This generated ∼4.5 million 100bp single end reads (median reads per sample = 51758) which were input into QIIME[50] for open reference OTU picking and taxonomic assignment using the UCLUST algorithm against the Greengenes (version 13_8) reference.

### Statistical analysis

A QIIME generated rarefied BIOM table (sampling depth 10,000 counts) was imported into R for statistical comparisons. The dataset was first filtered to remove OTUs that were only present at very low levels (max percent total community abundance less than 0.1% in all samples), or present in fewer than ∼10% of the samples. The resulting filtered dataset was used for downstream analysis. Statistical comparisons between groups were performed using non-parametric tests (e.g., Kruskal-Wallis tests) and multiple tests were corrected using q-value[51]. Tests with a q-value less than 0.2 and significant p-value (p < 0.05) were then subjected to pairwise Mann-Whitney U tests with Holm correction for multiple comparisons to determine which groups significantly differed. Fold changes in taxon abundance across groups were then calculated for each significant test. Here, fold change was defined as the quotient of the taxon abundance for an individual sample in the experimental group divided by the mean abundance of the normalizing group (i.e., the group to which a sample’s fold change is being compared). A small value (0.01 counts) was added to each observation in the OTU table prior to fold change analysis to prevent means of zero, which would produce spurious fold change values.

Indicator species analysis was used to identify OTUs that are characteristic of microbiomes from fish that were exposed to triclosan. Briefly, indicator species were identified for exposed (combined 7-day and 4-day exposure groups) and unexposed fish by using the labdsv R package. Q-values were calculated for each indicator species and poor indicators were removed (indicator values < 0.4, p-value > 0.05, or q-value > 0.2; labdsv::indval).

Alpha-diversity was measured using the Shannon index and statistical comparisons were calculated using Kruskal-Wallis tests that were subsequently subject to post-hoc pairwise Mann-Whitney U tests (Holm p-value correction) to determine group specific differences. Species richness was assessed using the rarefy function in the R package vegan (sampling depth 5,000 counts). Beta-diversity was measured using Bray-Curtis distance, and non-metric multidimensional scaling (NMDS) was used to quantify and visualize compositional similarity of communities. Significant differences in overall beta-diversity were calculated using analysis of similarity (ANOSIM; vegan::anosim), Permutational Multivariate Analysis of Variance (PERMANOVA, vegan::adonis), and environmental fit (vegan::envfit) using 5000 permutations for each test with the exception of envfit for which 10,000 permutations were calculated. Differences in Bray-Curtis dissimilarity between and within groups were calculated using Kruskal-Wallis tests and pairwise Mann-Whitney U tests (Holm correction).

### Microbial Correlation Network Analysis

Correlation networks of microbial abundance were constructed for each group (unexposed, 4-day, and 7-day) by calculating the Spearman’s rank correlation of OTU abundances. Weak (|rho| < 0.5) correlations and those that failed to reach significance (p > 0.05) and q-value thresholds (q > 0.2) were filtered. The remaining correlations were used to establish an interaction network where weighted nodes represent OTUs and edges represent the correlation coefficient calculated for a pair of OTUs. Next, networks were trimmed of self and duplicate edges before statistical analysis and visualization. Network statistics were calculated using the R package igraph and differences between exposure groups were tested using the Kruskall-Wallis and pairwise Mann-Whitney U tests with Holm correction. Pairwise Fisher’s exact tests were used to determine if proportions of negative and positive associations were different between unexposed and exposed groups. Pairwise p-values were adjusted using Holm’s method. Network community structure was detected using the fastgreedy.community function in the igraph R package.

## Results

### Triclosan exposure associated with shifts in microbial community structure

We established an experimental designed aimed at determining whether short-term, repeated exposure to triclosan can affect adult zebrafish gut microbial communities (Fig 1). Zebrafish were separated into three replicate tanks per treatment group (unexposed, four-day exposure, seven-day exposure; n=5 fish/tank). These fish were then fed diets that contained triclosan for four or seven days or diets without triclosan (control diet). After seven days the fish were euthanized and intestinal contents collected. We then constructed and sequenced 16S rRNA amplicon libraries from the intestinal contents samples.

**Fig 1.**
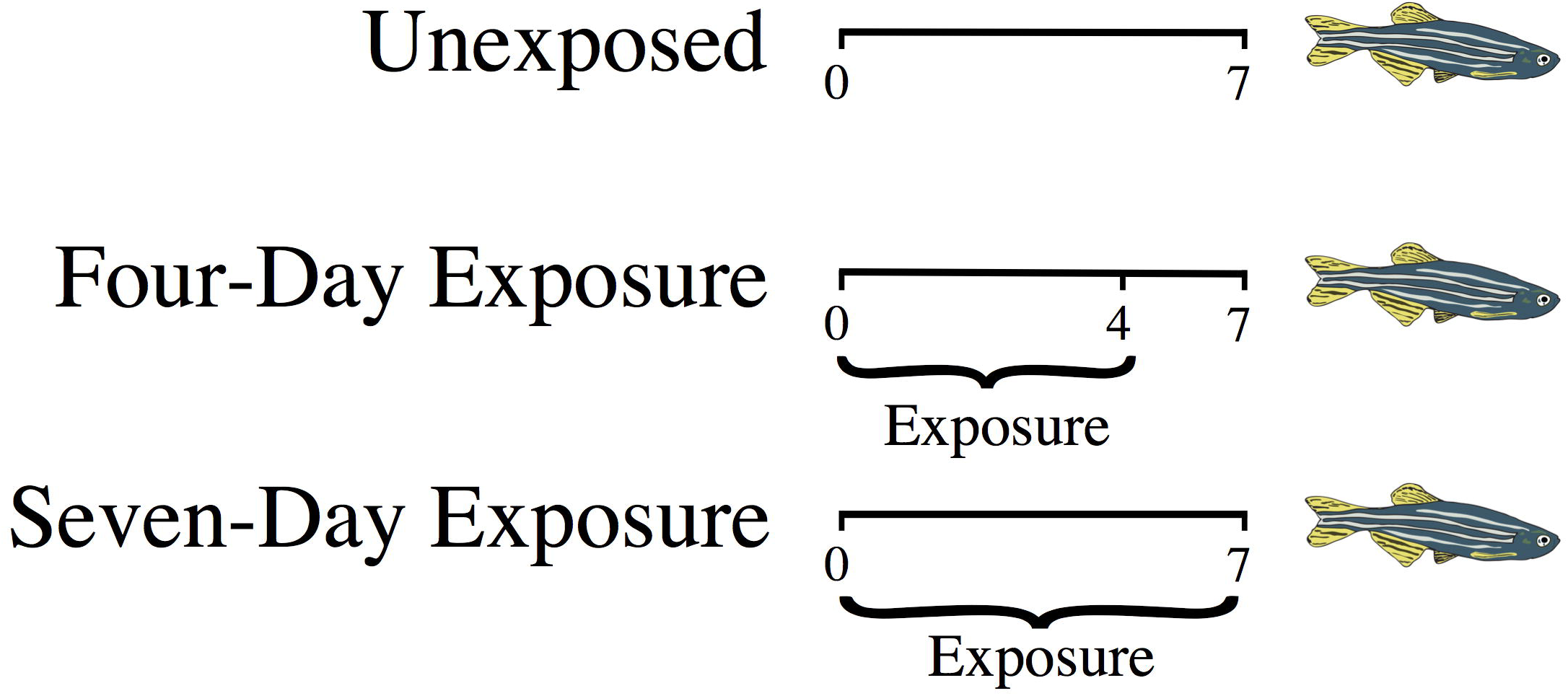
Triclosan exposure design. A schematic diagram of the experimental groups and triclosan exposures. Labeled tick marks represent days.

Consistent with previous work in zebrafish[34,52] and other aquatic organisms[53], Proteobacteria, and Fusobacteria, dominated the zebrafish gut microbiome (Fig 2A). At lower taxonomic levels the genera *Cetobacterium, Shewanella, Aeromonas,* the family Aeromonadaceae, and the class CK-1C4-19 were highly abundant in all groups. We then measured the beta-diversity among samples to quantify the impact of triclosan exposure on gut microbiome structure. Our analysis reveals a significant association (environmental fit p < 0.001; ANOSIM p < 0.05; PERMANOVA p < 0.05) between triclosan exposure and microbiome composition (Fig 2B). Additionally, there was a significant increase in intra-group Bray-Curtis dissimilarity between the unexposed and seven-day exposure groups (p < 0.05) indicating that the microbiomes of animals exposed to triclosan for 7 days were significantly more variable than unexposed animals (Fig 2C). We also observe differences in alpha-diversity among populations, finding that the seven-day exposure group has depreciated Shannon entropy relative to the four-day (p < 0.05) population (Fig 2D). Concordantly, species richness was reduced in the seven-day exposure animals relative to the four-day (p < 0.01). Taken together these data indicate that triclosan exposure results in destabilization and restructuring of microbial communities.

**Fig 2.**
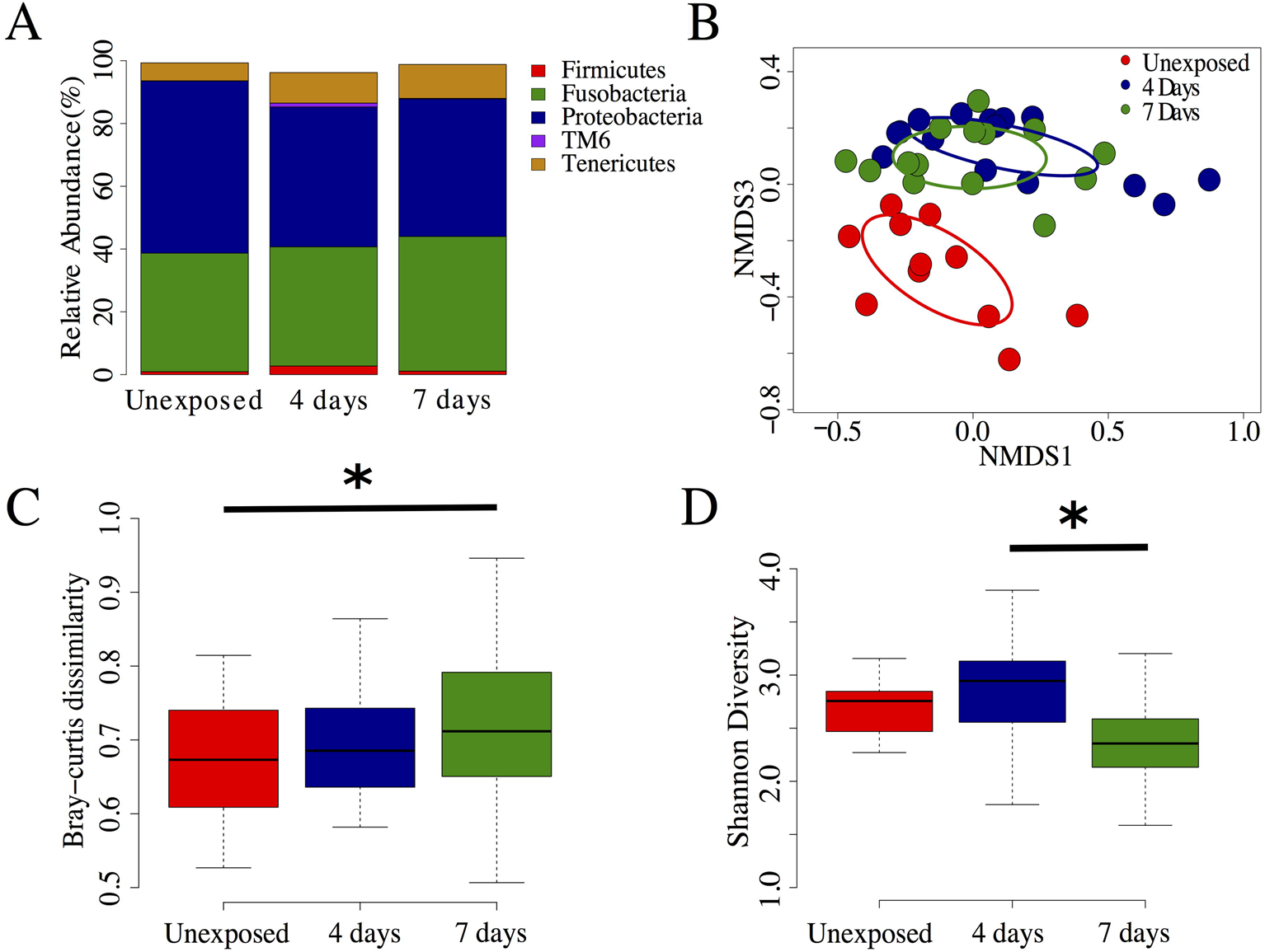
Triclosan exposure is associated with altered microbial community structure. (A) Phyla level taxa plot of most abundant taxa. (B) Non-metric multidimensional scaling analysis of unexposed (red dots), four-day (blue dots) and seven-day (green dots) exposure group’s microbial communities. Colored ellipses represent the 99.9% confidence interval for standard error of each group. (C) Comparisons of within group Bray-Curtis dissimilarity between groups. (D) Shannon diversity between exposure groups. Significant p-values (p < 0.05) are denoted with an asterisk.

### Zebrafish carry triclosan sensitive and resistant taxa

Triclosan inhibits bacterial growth by interfering with fatty acid synthesis through binding the enoyl-acyl carrier protein reductase enzyme (*Fabl*) [54]. However, several triclosan resistant mechanisms have been described and include mutations in triclosan target enzymes, increased enzyme expression, degradation of triclosan, and active efflux[55–57]. We reasoned that if resistant taxa exist in the microbiomes of zebrafish, their abundance should be increased or unchanged in exposed animals. Conversely, the abundance of susceptible microbes would be decreased. To determine if zebrafish microbiomes harbor resistant or susceptible microbes, we compared the abundance of OTUs and phylotypes between exposed and unexposed groups. Altered abundance was observed for 32 unique OTUs in the triclosan exposed fish when compared to unexposed fish. Triclosan exposure was indeed associated with changes in taxa that are consistent with the hypothesis that resistant and susceptible microbes are present in the zebrafish gut. For example, microbes phylotyped to the family Enterobacteriaceae were decreased in abundance in both exposure groups when compared to controls. These data were consistent with tests of OTU abundance, wherein a large portion of significantly altered OTUs in the exposed groups (67% in the four-and 57% in the seven-day exposure groups) were associated with the family Enterobacteriaceae (Fig 3 A,B). Notably all of these OTUs decreased in abundance in both exposure groups and many OTUs were associated with the genus *Plesiomonas.* Similarly, an OTU associated with the Aeromonadaceae family was also decreased in both exposure groups. These taxa appear to be more susceptible or less able to inhabit a gut niche environment associated with triclosan exposure. In addition, these taxa appear to have protracted vulnerability to triclosan exposure (i.e., taxa abundance does not recover to levels consistent with unexposed taxa abundance after removal of triclosan). In contrast, three OTUs associated the genera Chitinilyticum decreased in abundance in the seven-day exposure group when compared to unexposed controls, however, the abundance of these organisms was similar to unexposed controls in the four-day exposure group (Fig 3B). This potentially indicates that while these microbes are susceptible to triclosan exposure they are resilient to this perturbation (i.e., their abundance returned to levels of the control after removal of triclosan).

**Fig 3.**
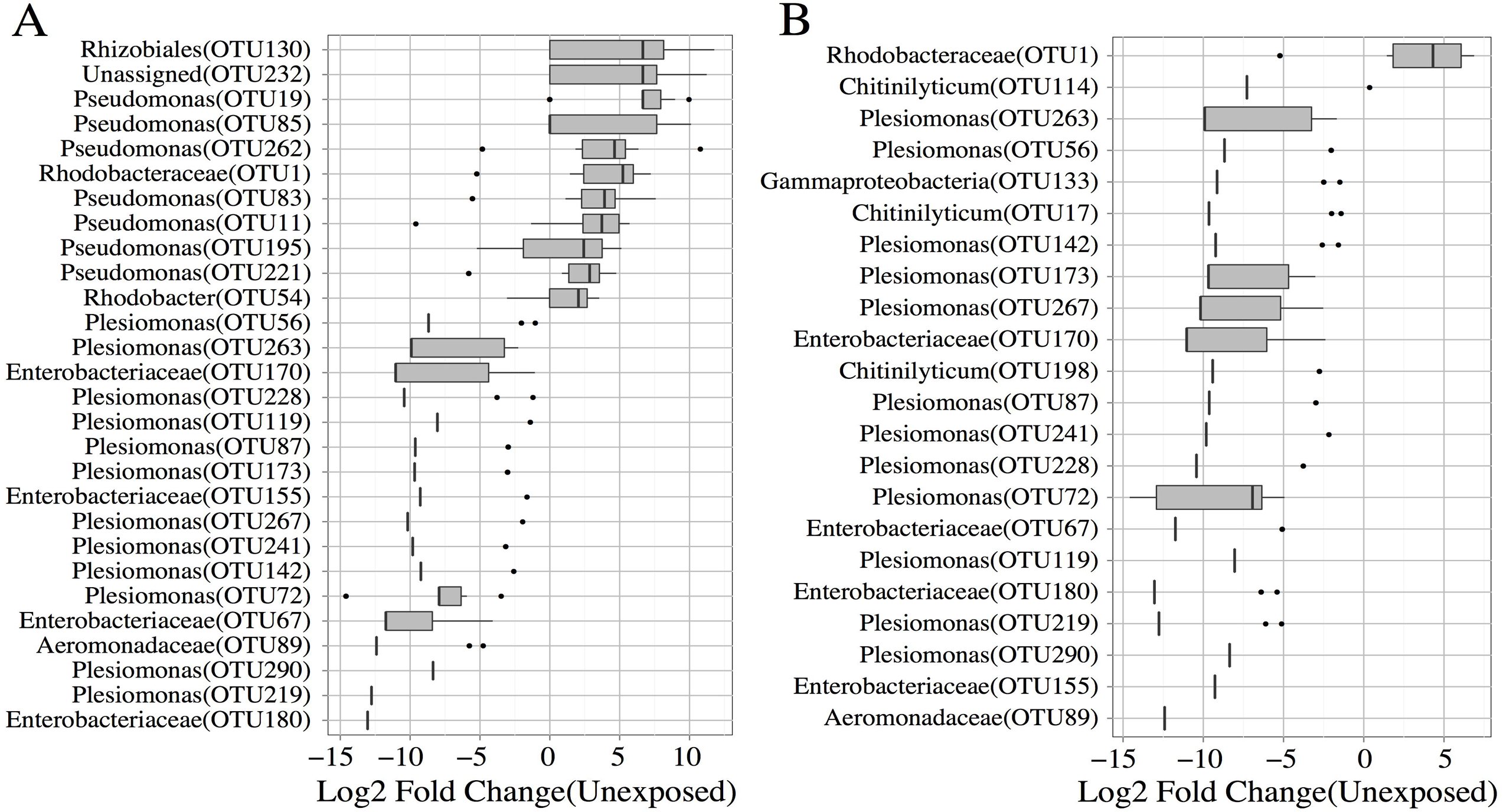
Triclosan exposure is associated with significant alterations in OTU abundance. Fold change values for OTUs that were significantly altered in abundance in the (A) four-day, and (B) seven day exposure groups. Genus level taxonomic assignments are provided and corresponding OTU IDs indicated inside parentheses.

We also identified microbes that are potentially resistant to triclosan in the zebrafish gut. We classified these organisms in two ways: (1) taxa that increased in abundance during exposure, and (2) taxa whose abundance was uninfluenced by triclosan exposure. A total of eleven OTUs increased in exposed groups when compared to controls. The majority of the OTUs that increased (∼64%) were associated with Pseudomonas and only significantly increased in the four-day exposure group (Fig 3A). Phylotype analysis confirmed that the abundance of the genus *Pseudomonas* was increased in the four-day group when compared to the unexposed groups. Similarly, an unknown member of Rhodobacteraceae family increased in abundance in the four-day exposure group. Concordantly, an OTU associated with this family was significantly increased in both exposure groups when compared to unexposed animals. Increased abundance of an OTU associated with the order Rhizobiales was also observed in the four-day exposure group. The increased abundance of these taxa indicates that they likely possess some degree of triclosan resistance or might have a higher relative fitness for inhabiting a triclosan exposed gut environment compared to other taxa. Indeed members of both the genus *Pseudomonas* and the family Rhodobacteraceae are known to be resistant to triclosan[58,59].

Finally, we examined taxa whose relative abundance was unaltered by triclosan exposure. We restricted this analysis to only highly abundant organisms (mean abundance > 100 counts) whose abundance did not change significantly during exposure (p > 0.05). This analysis identified sixteen OTUs, nine associated with the family Aeromonadaceae and six with the genus *Cetobacterium,* and one with the class CK-1C4-19 whose abundance remained unchanged despite triclosan exposure. Examination of the phylotype analysis confirmed that these taxa were unchanged during exposure. It is possible that these microbes encode triclosan resistance genes or that they constantly are replenished via environmental exposure (i.e., their water). Indeed triclosan resistant mutants carrying the FabV gene have been described for members of the family Aeromonadaceae [60]. Taken together these results indicate that the zebrafish gut harbors microbes both resistant and susceptible to triclosan exposure and that triclosan exposure is associated with altered abundance of specific taxa in this compartment.

### Triclosan exposed environments are associated with unique indicator taxa

One potential application of studying the effects of environmental chemicals on microbiome is that it might be possible to use shifts in these communities as biomarkers for these specific exposures. Ideally biomarkers are present at high abundance in only one group under investigation. However, the inferential techniques used to examine abundance do not consider presence and absence in their calculations. Thus, significant differences may exist between groups even if each group has relatively high abundance of a given OTU. This means that not all OTUs that differ significantly in abundance across groups are suitable biomarkers. To account for this we investigated whether any zebrafish gut microbiota produce predictive patterns of triclosan exposure duration through the use of indicator species analysis[61]. Importantly, this analysis considers both the abundance and frequency of occurrence (i.e., the number of samples in which a taxon is has an abundance greater than zero) of each taxon or OTU across samples, and assigns a value that indicates how representative each taxon or OTU is for each treatment group (i.e., an indicator value). We reasoned that this analysis could potentially identify biomarkers unique to specific environments or environmental chemical exposure.

We asked if there were any taxa that characterized triclosan-exposed environments and unexposed environments. For this analysis the seven-day and four-day exposure groups were combined and compared to the unexposed fish. We identified a total of 18 indicators of the unexposed group, all but two of which were associated with the family Enterobacteriaceae (Fig 4A), and seven indicators of triclosan exposed fish. These seven indicators are all OTUs associated with *Pseudomonas,* Rhodobacteraceae, Rhizobiales, and CK-1C4-19 (Fig 4B). Many of the indicators for exposed and unexposed environments overlapped with taxa we identified above as resistant and resilient. These results indicate that many, but not all, of the taxa whose abundance was significantly altered can be considered potential biomarkers for triclosan exposure.

**Fig 4.**
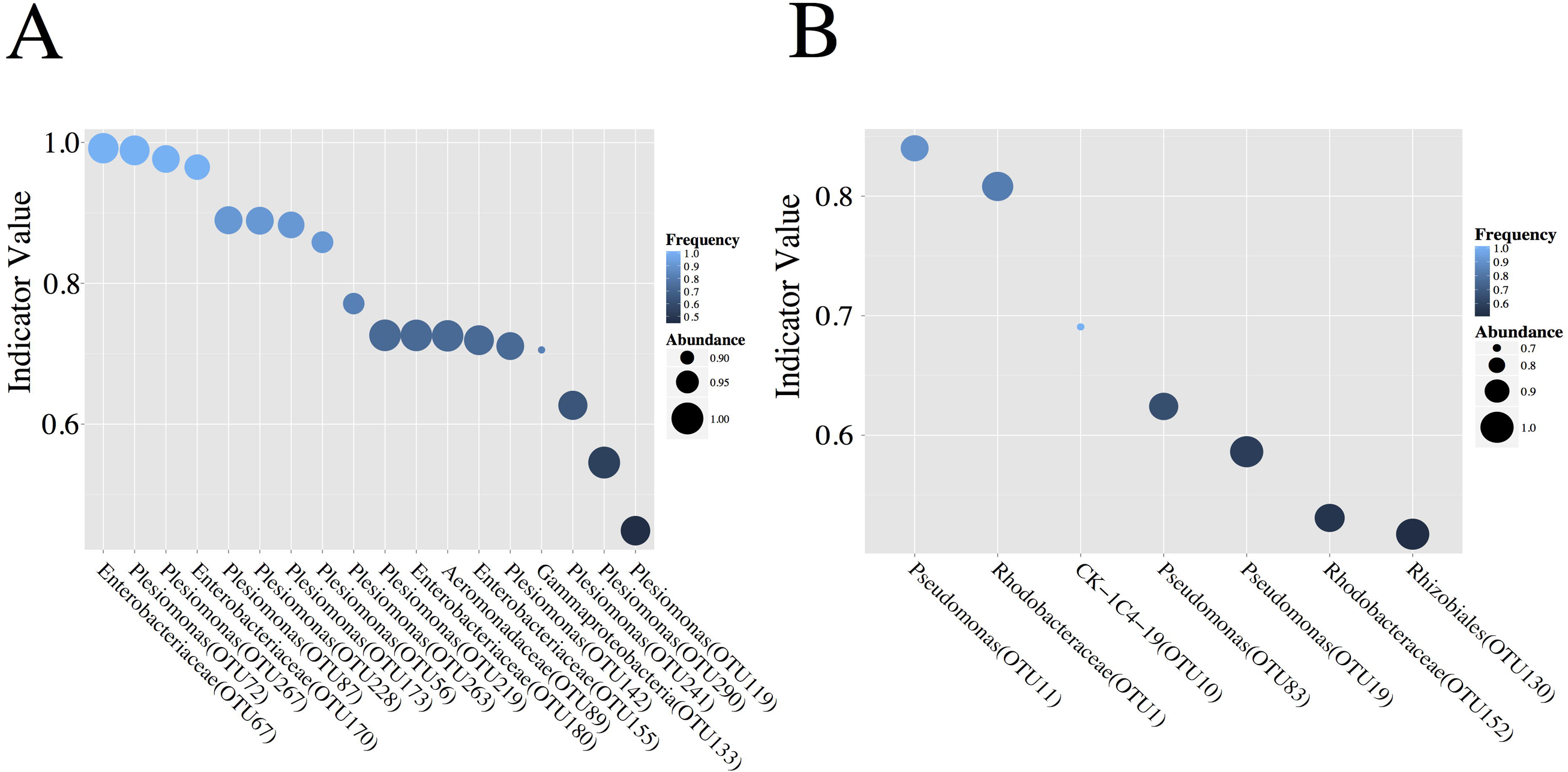
Triclosan exposure is associated with unique indicator OTUs. Significant indicator taxa for the (A) unexposed and (B) exposed fish. The size of each point is proportional to its class-wide relative abundance, and its color is proportional to its class-wide frequency.

### Altered microbial correlation network parameters is associate with triclosan exposure

The gastrointestinal ecosystem is diverse and complex, and the microbes that inhabit this space form intricate interactions with other microbes that are crucial to the operation of this ecosystem. While much has been learned about how perturbations to the gut microbiome impact its structure and diversity, less is known about how these perturbations impact the ecological interaction of the taxa that comprise the microbiome. We used inferential techniques[62,63] and comparative network topological analysis to assess whether triclosan exposure perturbs how gut microbiota ecologically relate to one another.

Correlations of abundance were calculated for all pairs of microbes separately for each group and these correlation datasets were filtered to remove weak interactions and those that did not reach significance or pass q-value filtering (see methods). Those pairs that passed our criteria were then used to create correlation networks. After filtering, the networks were comprised of 195, 269, and 231 vertices for the unexposed, four-, and seven-day groups respectively (Fig 5 A-C). Interestingly, the majority of these vertices were shared between the three groups (Fig 5D), however, triclosan exposed fish had starkly different topological parameters and a significant enrichment for negative correlations between taxa (p < 1x10^-6^). The increase in negative correlations was accompanied by an overall increase in degree centrality (number of edges per vertex) of both the four-day exposure group (p < 1.0x10^-12^) and the seven-day exposure group (p < 0.0005) when compared to the unexposed groups (Fig 5E). The increase in degree distribution was robust to directionality of the edge (i.e., there was increased numbers of both positive and negative distributions). Moreover, the proportion of negative edges per vertex increased in the exposed groups, as did the proportion of nodes that contained at least one negative edge (Table 1). The enrichment for negative correlations might indicate increased competition among intestinal microbes for niche space or nutrient availability.

**Fig 5.**
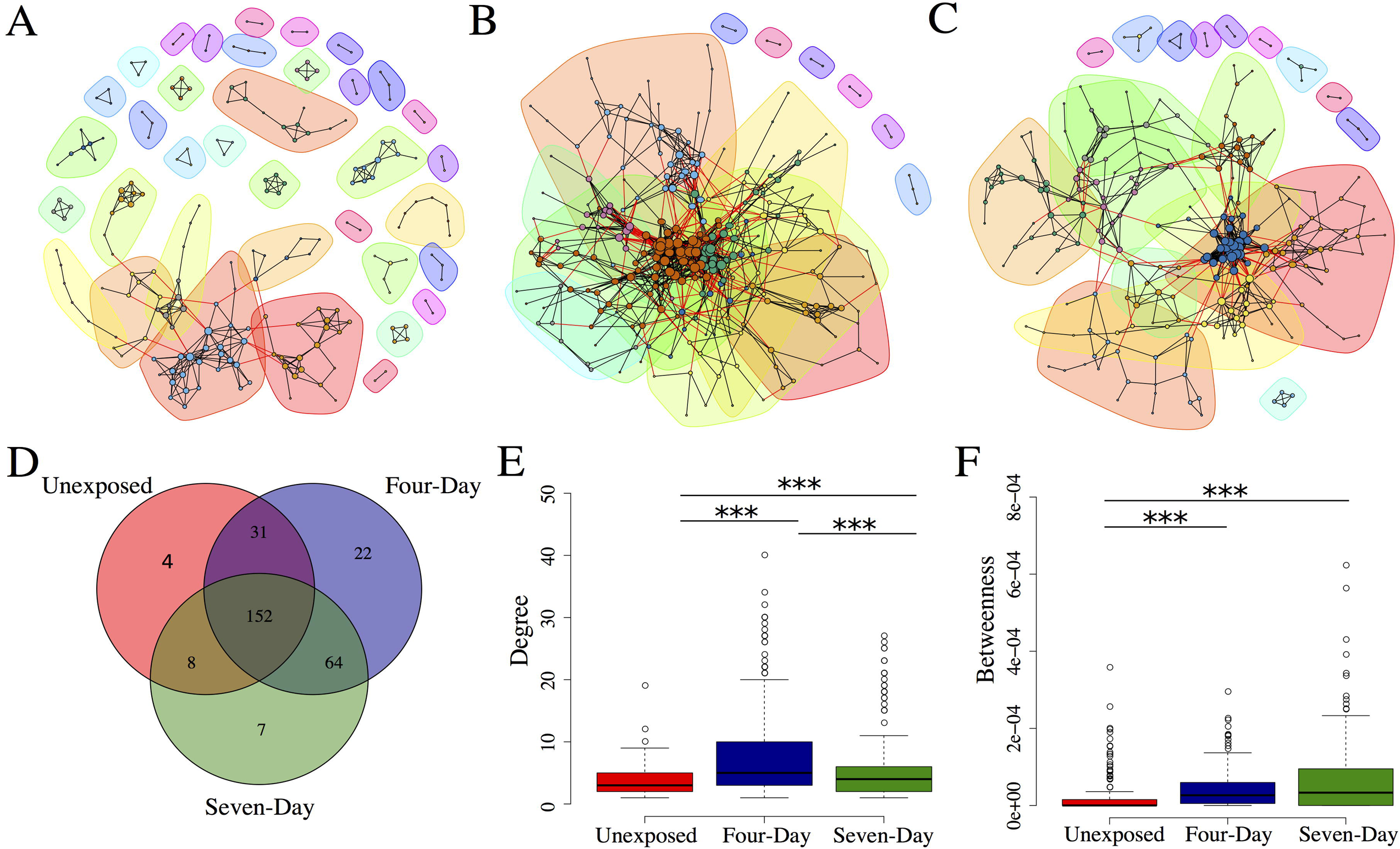
Triclosan exposure is associated with alterations in microbial correlation networks. Interaction networks for microbial communities in (A) unexposed, (B) four-day exposure and (C) seven-day exposure groups. Communities (subgraphs) are colored by identity. Each line represents an abundance correlation between two OTUs. The size of each vertex is proportional to its degree and each vertex is colored by its community identity. Edges between that connect a vertex from one community to a vertex of another are colored red. (D) Venn diagram of shared vertices between networks. (E) Degree and (F) betweenness centrality distribution for all vertices in network. *** p < 0.001.

**Table 1.** Network Properties.

| Group | Vertices | Total Edges | Positive Edges | Negative Edges | Mean Degree | Mean Betweenness | Vertices > 1 negative edge | %Vertices > 1 negative edge | Vertices > 1 positive edge | %Vertices > 1 positive edge | Communities |
| --- | --- | --- | --- | --- | --- | --- | --- | --- | --- | --- | --- |
| Unexposed | 195 | 324 | 289 | 35 | 3.3 | $2.2 \times 10^{-5}$ | 43 | 22.1 | 189 | 96.9 | 36 |
| Four-Day | 269 | 1040 | 792 | 248 | 7.7 | $4 \times 10^{-5}$ | 155 | 57.6 | 258 | 95.9 | 14 |
| Seven-Day | 231 | 627 | 460 | 167 | 5.4 | $6.2 \times 10^{-5}$ | 102 | 44.2 | 221 | 95.7 | 19 |

We then asked how exposure to triclosan influenced the connectivity of networks by measuring betweenness centrality. Betweenness measures how many shortest paths between every pair of vertices pass through a specific vertex and represents an approximation of a vertex’s influence on information flow through a network[64]. The networks from triclosan-exposed fish had increased betweenness centrality when compared to unexposed fish (p < 1.0x10-^10^both groups; Fig 5F). This was consistent with the increased degree distribution in these groups and indicates that the vertices in correlation networks of exposed fish tend to have a greater influence and a higher degree of connectivity than do unexposed networks. This increased connectivity was also reflected in a decreased number of communities in the networks of triclosan-exposed fish (Fig 5A-C; Table 1). Community detection analysis attempts to divide a network into communities of vertices that share many edges with members of the community but few with vertices outside the community[64] and can be used to infer relationships between its members[65–67]. Together these data indicate that triclosan exposure is associated with topological rearrangement of microbial correlation networks and that these changes are manifested primarily as increased connectivity in exposed communities.

## Discussion

A growing body of evidence suggests that triclosan might alter host physiology [22,42], disrupt environmental microbial communities[45,46], and increase antimicrobial and antibiotic resistance in the environment and in laboratory bacterial strains[38,68]. Narrowe *et al.* recently demonstrated that triclosan exposure alters the microbiome of juvenile fathead minnows *(Pimephales promelas)*[47]. However, they did not specifically investigate the impact of triclosan exposure on adults, which are known to be more refractory to microbiome perturbation[69]. In the present study we have investigated the impact of triclosan on the gut microbiomes of adult zebrafish using a dietary triclosan exposure regimen. Similar to Narrowe *et al.,* we find that triclosan exposure is associated with disruption of the composition of microbial communities of adult fish over short time intervals. Our results complement and extend the findings of Narrowe et al. in several ways. For example, despite examining different life stages (i.e., adult vs. juvenile) both studies identified similar taxa that change as a result of triclosan exposure (e.g., *Rhodobacter, Pseudomonas,* etc.). These data bolster the strength of both studies and indicate that the effects of triclosan exposure are, at least in part, robust to developmental stage of the organism, and that there may be conserved patterns of microbiome sensitivity across host species. Through additional analyses we were also able to identify potential biomarkers of triclosan exposure and evidence indicates that triclosan exposure perturbs how gut microbiota interact. This study provides novels insights into the shifts in microbial communities and their interactions that are associated with triclosan exposure.

Antibiotics are strong perturbing agents of the microbiome and are associated with dramatic[69], and in some cases long lasting[10,11], alterations of the microbial community composition. Microbial community disruption by antibiotics can lead to dysbiosis and potentially to increased susceptibility to infection with opportunistic pathogens. For example, disruption of the gut microbiome by antibiotics contributes to the colonization efficiency *Clostridium difficile* in humans and mice[70,71]. Moreover, the microbial community compositions that confer resistance to *C. difficile* are varied and no one taxon confers complete protection. Triclosan exposure was associated with a similar, but more modest, restructuring of community composition when compared to clinical antibiotics. For example, although there were clear differences in beta-diversity between exposed and unexposed fish relatively few distinct taxa that changed during exposure. However, the changes in taxon abundance we did observe were consistent with the hypothesis that triclosan resistant taxa would accumulate in the exposed environment. Previous studies have found that concentrations of triclosan in sediment are correlated with the number of triclosan resistant bacteria in these communities [72]. Although we did not quantify triclosan resistant bacteria directly in this study, it is well established that many members of the genus *Pseudomonas,* which increased in the present study, are highly resistant to triclosan[56,58,73]. Mutants with increased resistance to triclosan have also been described for members of Rhodobacteraceae[59]. Conversely many members of the family Enterobacteriaceae are known to be susceptible to triclosan [74,75], although resistant mutants have also been described[76]. These observations raise the possibility that triclosan exposure might lead to increased abundance of triclosan resistant organisms in the gut. This is potentially concerning as triclosan exposure has also been associated with increased abundance and diversity of antimicrobial genes in laboratory strains[68]. Recent evidence also suggests that some mechanisms of triclosan resistance might be able to be transferred horizontally[77]. Future studies should endeavor to clarify if OTUs that increase during exposure possess adaptations that confer triclosan resistance and if triclosan exposure is correlated with increased diversity of antibiotic resistance genes in microbial communities.

Indicator species analysis identified several gut microbiota that statistically stratify exposed and unexposed fish and may serve as biomarkers of exposure. Specially, triclosan exposed fish have low abundance of several OTUs associated with the family Enterobacteriaceae, and increased abundance of OTUs associated with Pseudomonas. Characterization and cataloging of indicator species of environmental chemical exposure in this manner may uncover biomarkers that could be useful in environmental and health monitoring. For example, samples obtained from aquatic organisms could be screened against a catalog of known environmental contaminants with defined indicator species to determine to presence or identity of a toxicant. Alternatively, clinical samples could be screened against a database to determine if a patient had any known chemical exposure. Indicator species analysis of microbial communities has previously been used to identify biomarkers of mucosal disease[78,79] highlighting the potential of this technique.

Notably, triclosan exposure resulted in substantial topological changes to microbial correlation networks, indicating that exposure might alter how microbes interact and communicate with one another in the gut. Unexpectedly, the mean degree, and betweenness of the networks increased with triclosan exposure indicating more connective networks. Increased degree and betweenness could be, in part, explained by succession in these communities. For example, in the unexposed gut environment there likely exist a number of metabolic and physiological interactions between microbes, however when this environment is perturbed and the abundance of microbes shift, these relationships breakdown and niche space is opened. As new triclosan resistant microbes begin to immigrate into the niche space vacated by triclosan sensitive microbes new metabolic and physiological partnerships form. Alternatively, the increased interactions might simply reflect similar growth kinetics of species immigrating to these vacant niches. Regardless, increased positive correlations in the exposed networks suggests increased cooperative interactions between microbes in these networks. Although counter-intuitive, this increase may indicate that these networks are less, not more, stable as cooperation may create dependencies between microbes[80]. The increased number of negative correlations in the exposed communities might indicate increased competition for nutrient or niche space in exposed communities. While competition can be stabilizing[80] it can also produce undesired effects such as loss of taxa beneficial to the host. Importantly, changes in the interaction landscape among microbiota may affect host physiology. For example, children that develop type-1 diabetes had significant differences in the microbial interaction networks at a young age[63]. Future research should explore the physiological outcomes of the altered network topology associated with triclosan exposure in zebrafish.

Given the frequency and diversity of environmental chemical exposure to which animals are subject and the importance of the gut microbiome to animal health, understanding what the impacts of these exposures are on the microbiome is of paramount importance. Although the impacts of acute exposures may be modest, some chemical exposures might also have cumulative impacts. Therefore it is not only important to understand the individual effects of these toxicants, but also the synergistic impacts. We posit that zebrafish lends itself well to studies of this kind as large samples sizes are more manageable and economically feasible than in other animal models. As seen here, access to a large number of individuals enables the resolution of statistically subtle, but biologically meaningful effects and affords the power needed to understand how microbial correlation networks are affected by exposure. Additionally, these features of the zebrafish model mean that a large array of chemicals across a spectrum of concentrations can be screened. Chemicals identified as potentially important perturbing agents of the microbiome can then be further examined in other animal models or in epidemiological investigations in humans. Moreover, the results of the present study are largely consistent with the findings of Narrowe *et al.* indicating that the effects of chemical exposure may be a conserved feature of the microbiome across fish species. This observation, coupled with the similarity between the microbiomes of zebrafish and other aquatic organisms[53], indicates that the zebrafish could be used to model the effects of environmental chemical exposure on the microbiome and health of wild and farm-raised fish. We used this model to demonstrate that triclosan exposure is associated with restructuring of gut microbial communities and altered topology of microbial correlation networks in adult fish. One caveat of this study is that fish were exposed to triclosan through their diet, which means that the dose that an individual receives varies with the amount of food it consumes. Future work should investigate how the route and concentration of exposure impacts the zebrafish microbiome, and quantify how these perturbations impact zebrafish physiology. Regardless, these data add to a growing body of evidence that indicates that triclosan exposure might impact hosts in previously unexpected the ways and underscore the utility of zebrafish as an experimental tool for understanding how environmental chemical exposure impacts the gut microbiome.

## Acknowledgements

The authors thank the staff of SARL for the assistance in animal studies and to the members of the Sharpton lab for their helpful discussion of the manuscript. We also thank members of the CGRB for their assistance in sequencing libraries and in assistance with the computational infrastructure.

## References

1. Bohnhoff M, Drake BL, Miller CP. Effect of streptomycin on susceptibility of intestinal tract to experimental Salmonella infection. Exp Biol Med. SAGE Publications; 1954;86: 132–137.

2. van der Waaij D, Berghuis-de Vries JM, Lekkerkerk Lekkerkerk-v. Colonization resistance of the digestive tract in conventional and antibiotic-treated mice. J Hyg (Lond). 1971;69: 405–411. doi:10.1017/S0022172400021653

3. Bouskra D, Brézillon C, Bérard M, Werts C, Varona R, Boneca IG, et al. Lymphoid tissue genesis induced by commensals through NOD1 regulates intestinal homeostasis. Nature. 2008;456: 507–510. doi:10.1038/nature07450

4. Mazmanian SK, Round JL, Kasper DL. A microbial symbiosis factor prevents intestinal inflammatory disease. Nature. 2008;453: 620–625. doi:10.1038/nature07008

5. Donohoe DR, Wali A, Brylawski BP, Bultman SJ. Microbial Regulation of Glucose Metabolism and Cell-Cycle Progression in Mammalian Colonocytes. PLoS One. 2012;7. doi:10.1371/journal.pone.0046589

6. Qin J, Li Y, Cai Z, Li S, Zhu J, Zhang F, et al. A metagenome-wide association study of gut microbiota in type 2 diabetes. Nature. 2012. pp. 55–60. doi:10.1038/nature11450

7. Koeth R a, Wang Z, Levison BS, Buffa J a, Org E, Sheehy BT, et al. Intestinal microbiota metabolism of L-carnitine, a nutrient in red meat, promotes atherosclerosis. Nat Med. 2013;19: 576–85. doi:10.1038/nm.3145

8. Scher JU, Sczesnak A, Longman RS, Segata N, Ubeda C, Bielski C, et al. Expansion of intestinal Prevotella copri correlates with enhanced susceptibility to arthritis. Elife. 2013;2013.doi:10.7554/eLife.01202.001

9. Smith MI, Yatsunenko T, Manary MJ, Trehan I, Mkakosya R, Cheng J, et al. Gut microbiomes of Malawian twin pairs discordant for kwashiorkor. Science. 2013;339: 548–54. doi:10.1126/science.1229000

10. Dethlefsen L, Relman DA. Incomplete recovery and individualized responses of the human distal gut microbiota to repeated antibiotic perturbation. Proc Natl Acad Sci U S A. 2011;108 Suppl: 4554–4561. doi:10.1073/pnas.1000087107

11. Jernberg C, Löfmark S, Edlund C, Jansson JK. Long-term ecological impacts of antibiotic administration on the human intestinal microbiota. ISME J. 2007;1: 56–66. doi:10.1038/ismej.2007.3

12. Glavan TW, Gaulke CA, Santos Rocha C, Sankaran-Walters S, Hirao LA, Raffatellu M, et al. Gut immune dysfunction through impaired innate pattern recognition receptor expression and gut microbiota dysbiosis in chronic SIV infection. Mucosal Immunol. Society for Mucosal Immunology; 2015; Available: http://dx.doi.org/10.1038/mi.2015.92

13. Nelson AM, Walk ST, Taube S, Taniuchi M, Houpt ER, Wobus CE, et al. Disruption of the human gut microbiota following Norovirus infection. PLoS One. 2012;7: e48224. doi:10.1371/journal.pone.0048224

14. Muegge BD, Kuczynski J, Knights D, Clemente JC, González A, Fontana L, et al. Diet drives convergence in gut microbiome functions across mammalian phylogeny and within humans. Science. 2011;332: 970–974. doi:10.1126/science.1198719

15. David LA, Maurice CF, Carmody RN, Gootenberg DB, Button JE, Wolfe BE, et al. Diet rapidly and reproducibly alters the human gut microbiome. Nature. 2014;505: 559–63. doi:10.1038/nature12820

16. Rappaport SM, Barupal DK, Wishart D, Vineis P, Scalbert A. The blood exposome and its role in discovering causes of disease. Environ Health Perspect. 2014; 122:769–774. doi:10.1289/ehp.1308015

17. Wild CP, Scalbert A, Herceg Z. Measuring the exposome: a powerful basis for evaluating environmental exposures and cancer risk. Environ Mol Mutagen. 2013;54: 480–99. doi:10.1002/em.21777

18. Nicholson JK, Holmes E, Kinross J, Burcelin R, Gibson G, Jia W, et al. Host-gut microbiota metabolic interactions. Science (80-). 2012;336: 1262–7. doi:10.1126/science.1223813

19. Lu K, Abo RP, Schlieper KA, Graffam ME, Levine S, Wishnok JS, et al. Arsenic exposure perturbs the gut microbiome and its metabolic profile in mice: An integrated metagenomics and metabolomics analysis. Environ Health Perspect. 2014;122: 284–291. doi:10.1289/ehp.1307429

20. Breton J, Massart S, Vandamme P, De Brandt E, Pot B, Foligné B. Ecotoxicology inside the gut: impact of heavy metals on the mouse microbiome. BMC Pharmacol Toxicol. 2013;14: 62. doi:10.1186/2050-6511-14-62

21. Liu Y, Li Y, Liu K, Shen J. Exposing to cadmium stress cause profound toxic effect on microbiota of the mice intestinal tract. PLoS One. 2014;9. doi:10.1371/journal.pone.0085323

22. Pinto PIS, Guerreiro EM, Power DM. Triclosan interferes with the thyroid axis in the zebrafish (Danio rerio). Toxicology Research. 2013. doi:10.1039/c2tx20005h

23. Darbre PD, Harvey PW. Paraben esters: Review of recent studies of endocrine toxicity, absorption, esterase and human exposure, and discussion of potential human health risks. Journal of Applied Toxicology. 2008. pp. 561–578. doi:10.1002/jat.1358

24. Magee EA, Edmond LM, Tasker SM, Kong SC, Curno R, Cummings JH. Associations between diet and disease activity in ulcerative colitis patients using a novel method of data analysis. Nutr J. 2005;4: 7. doi:10.1186/1475-2891-4-7

25. Spitsbergen J, Kent M. The State of the Art of the Zebrafish Model for Toxicology and Toxicologic Pathology Research - Advantages and Current Limitations. Toxicol Pathol. 2003;31: 62–87. doi:10.1080/01926230390174959

26. Kent ML, Buchner C, Barton C, Tanguay RL. Toxicity of chlorine to zebrafish embryos. Dis Aquat Organ. 2014;107: 235–40. doi:10.3354/dao02683

27. Truong L, Denardo MA, Kundu S, Collins TJ, Tanguay RL. Zebrafish Assays as Developmental Toxicity Indicators in The Design of TAML Oxidation Catalysts. Green Chem. 2013;15: 2339–2343. doi:10.1039/C3GC40376A

28. Wienholds E, van Eeden F, Kosters M, Mudde J, Plasterk RH a, Cuppen E. Efficient target-selected mutagenesis in zebrafish. Genome Res. 2003;13: 2700–2707. doi:10.1101/gr.1725103

29. Grunwald DJ, Eisen JS. Headwaters of the zebrafish — emergence of a new model vertebrate. Nat Rev Genet. 2002;3: 717–724. doi:10.1038/nrg891

30. Burns CG, Milan DJ, Grande EJ, Rottbauer W, MacRae C a, Fishman MC. High-throughput assay for small molecules that modulate zebrafish embryonic heart rate. Nat Chem Biol. 2005;1: 263–264. doi:10.1038/nchembio732

31. Truong L, Mandrell D, Mandrell R, Simonich M, Tanguay RL. A rapid throughput approach identifies cognitive deficits in adult zebrafish from developmental exposure to polybrominated flame retardants. Neurotoxicology. 2014;43: 134–42. doi:10.1016/j.neuro.2014.03.005

32. Stephens WZ, Wiles TJ, Martinez ES, Jemielita M, Burns AR, Parthasarathy R, et al. Identification of Population Bottlenecks and Colonization Factors during Assembly of Bacterial Communities within the Zebrafish Intestine. MBio. 2015;6: e01163–15–. doi:10.1128/mBio.01163-15

33. Jemielita M, Taormina MJ, Burns AR, Hampton JS, Rolig AS, Guillemin K, et al. Spatial and Temporal Features of the Growth of a Bacterial Species Colonizing the Zebrafish Gut. MBio. 2014;5: 1–8. doi:10.1128/mBio.01751-14

34. Zac Stephens W, Burns AR, Stagaman K, Wong S, Rawls JF, Guillemin K, et al. The composition of the zebrafish intestinal microbial community varies across development. ISME J. International Society for Microbial Ecology; 2015; Available: http://dx.doi.org/10.1038/ismej.2015.140

35. Rolig AS, Parthasarathy R, Burns AR, Bohannan BJM, Guillemin K. Individual Members of the Microbiota Disproportionately Modulate Host Innate Immune Responses. Cell Host Microbe. Elsevier; 2015;18: 613–620.

36. Kanther M, Tomkovich S, Xiaolun S, Grosser MR, Koo J, Flynn EJ, et al. Commensal microbiota stimulate systemic neutrophil migration through induction of Serum amyloid A. Cell Microbiol. 2014;16: 1053–1067. doi:10.1111/cmi.12257

37. Semova I, Carten JD, Stombaugh J, Mackey LC, Knight R, Farber SA, et al. Microbiota regulate intestinal absorption and metabolism of fatty acids in the zebrafish. Cell Host Microbe. 2012;12: 277–88. doi:10.1016/j.chom.2012.08.003

38. Schweizer HP. Triclosan: A widely used biocide and its link to antibiotics. FEMS Microbiology Letters. 2001. pp. 1–7. doi:10.1016/S0378-1097(01)00273-7

39. Sandborgh-Englund G, Adolfsson-Erici M, Odham G, Ekstrand J. Pharmacokinetics of triclosan following oral ingestion in humans. Journal of toxicology and environmental health. Part A. 2006. doi:10.1080/15287390600631706

40. Moss T, Howes D, Williams FM. Percutaneous penetration and dermal metabolism of triclosan (2,4, 4’-trichloro-2′-hydroxydiphenyl ether). Food Chem Toxicol. 2000;38: 361–370.

41. Allmyr M, Adolfsson-Erici M, McLachlan MS, Sandborgh-Englund G. Triclosan in plasma and milk from Swedish nursing mothers and their exposure via personal care products. Sci Total Environ. 2006;372: 87–93. doi:10.1016/j.scitotenv.2006.08.007

42. Stoker TE, Gibson EK, Zorrilla LM. Triclosan exposure modulates estrogen-dependent responses in the female wistar rat. Toxicol Sci. 2010;117: 45–53. doi:10.1093/toxsci/kfq180

43. Yueh M-F, Taniguchi K, Chen S, Evans RM, Hammock BD, Karin M, et al. The commonly used antimicrobial additive triclosan is a liver tumor promoter. Proc Natl Acad Sci U S A. 2014; doi:10.1073/pnas.1419119111

44. Wallet MA, Calderon NL, Alonso TR, Choe CS, Catalfamo DL, Lalane CJ, et al. Triclosan alters antimicrobial and inflammatory responses of epithelial cells. Oral Dis. 2013;19: 296–302. doi:10.1111/odi.12001

45. Harrow DI, Felker JM, Baker KH. Impacts of triclosan in greywater on soil microorganisms. Appl Environ Soil Sci. Hindawi Publishing Corporation; 2011;2011.

46. Nietch CT, Quinlan EL, Lazorchak JM, Impellitteri CA, Raikow D, Walters D. Effects of a chronic lower range of triclosan exposure on a stream mesocosm community. Environ Toxicol Chem. 2013;32: 2874–2887. doi:10.1002/etc.2385

47. Narrowe AB, Albuthi-Lantz M, Smith EP, Bower KJ, Roane TM, Vajda AM, et al. Perturbation and restoration of the fathead minnow gut microbiome after low-level triclosan exposure. Microbiome. BioMed Central Ltd; 2015;3: 6.

48. Caporaso JG, Lauber CL, Walters WA, Berg-Lyons D, Huntley J, Fierer N, et al. Ultra-high-throughput microbial community analysis on the Illumina HiSeq and MiSeq platforms. The ISME Journal. 2012. pp. 1621–1624. doi:10.1038/ismej.2012.8

49. Caporaso JG, Lauber CL, Walters WA, Berg-Lyons D, Lozupone CA, Turnbaugh PJ, et al. Global patterns of 16S rRNA diversity at a depth of millions of sequences per sample. Proc Natl Acad Sci U S A. 2011;108 Suppl: 4516–4522. doi:10.1073/pnas.1000080107

50. Caporaso JG, Kuczynski J, Stombaugh J, Bittinger K, Bushman FD, Costello EK, et al. QIIME allows analysis of high-throughput community sequencing data. Nature methods. 2010. pp. 335–336. doi:10.1038/nmeth.f.303

51. Storey JD. A direct approach to false discovery rates. J R Stat Soc Ser B (Statistical Methodol. Wiley Online Library; 2002;64: 479–498.

52. Roeselers G, Mittge EK, Stephens WZ, Parichy DM, Cavanaugh CM, Guillemin K, et al. Evidence for a core gut microbiota in the zebrafish. The ISME Journal. 2011. pp. 1595–1608. doi:10.1038/ismej.2011.38

53. Llewellyn MS, Boutin S, Hoseinifar SH, Derome N. Teleost microbiomes: the state of the art in their characterization, manipulation and importance in aquaculture and fisheries. Front Microbiol. 2014;5: 207. doi:10.3389/fmicb.2014.00207

54. Heath RJ, Rubin JR, Holland DR, Zhang E, Snow ME, Rock CO. Mechanism of triclosan inhibition of bacterial fatty acid synthesis. J Biol Chem. 1999;274: 11110–11114. doi:10.1074/jbc.274.16.11110

55. Kim YM, Murugesan K, Schmidt S, Bokare V, Jeon JR, Kim EJ, et al. Triclosan susceptibility and co-metabolism - A comparison for three aerobic pollutant-degrading bacteria. Bioresour Technol. 2011;102: 2206–2212. doi:10.1016/j.biortech.2010.10.009

56. Chuanchuen R, Beinlich K, Hoang TT, Becher A, Karkhoff-Schweizer RR, Schweizer HP. Cross-resistance between triclosan and antibiotics in Pseudomonas aeruginosa is mediated by multidrug efflux pumps: Exposure of a susceptible mutant strain to triclosan selects nfxB mutants overexpressing MexCD-OprJ. Antimicrob Agents Chemother. 2001;45: 428–432. doi:10.1128/AAC.45.2.428-432.2001

57. McMurry LM, Oethinger M, Levy SB. Triclosan targets lipid synthesis. Nature. 1998. pp. 531–532. doi:10.1038/28970

58. Zhu L, Lin J, Ma J, Cronan JE, Wang H. Triclosan resistance of Pseudomonas aeruginosa PAO1 is due to FabV, a triclosan-resistant enoyl-acyl carrier protein reductase. Antimicrob Agents Chemother. 2010;54: 689–698. doi:10.1128/AAC.01152-09

59. Lee IH, Kim EJ, Cho YH, Lee JK. Characterization of a novel enoyl-acyl carrier protein reductase of diazaborine-resistant Rhodobacter sphaeroides mutant. Biochem Biophys Res Commun. 2002;299: 621–627. doi:10.1016/S0006-291X(02)02702-X

60. Khan R, Lee MH, Joo H, Jung Y-H, Ahmad S, Choi J, et al. Triclosan Resistance in a Bacterial Fish Pathogen Aeromonas salmonicida subsp. salmonicida is Mediated by an Enoyl Reductase FabV. J Microbiol Biotechnol. 2015;25: 511–520. doi:10.4014/jmb.1407.07021

61. Dufreňe M, Legendre P. Species assemblages and indicator species: the need for a flexible asymmetrical approach. Ecol Monogr. 1997;67: 345–366. doi:10.1890/0012-9615(1997)067[0345:SAAIST]2.0.CO;2

62. Arumugam M, Raes J, Pelletier E, Le Paslier D, Yamada T, Mende DR, et al. Enterotypes of the human gut microbiome. Nature. 2011;473: 174–180. doi:10.1038/nature10187

63. Endesfelder D, Castell WZ, Ardissone A, Davis-Richardson AG, Achenbach P, Hagen M, et al. Compromised gut microbiota networks in children with anti-islet cell autoimmunity. Diabetes. 2014;63: 2006–2014. doi:10.2337/db13-1676

64. Newman M. Networks an Introduction [Internet]. Book. Oxford: Oxford University Press; 2010. doi:10.1093/acprof:oso/9780199206650.001.0001

65. Wilkinson DM, Huberman B a. A method for finding communities of related genes. Proc Natl Acad Sci U S A. 2004;101 Suppl: 5241–5248. doi:10.1073/pnas.0307740100

66. Rives AW, Galitski T. Modular organization of cellular networks. Proc Natl Acad Sci USA. 2003;100: 1128–1133. doi:10.1073/pnas.0237338100

67. Lewis ACF, Jones NS, Porter MA, Deane CM. The function of communities in protein interaction networks at multiple scales. BMC Syst Biol. 2010;4: 100. doi:10.1186/1752-0509-4-100

68. Yazdankhah SP, Scheie AA, Høiby EA, Lunestad B-T, Heir E, Fotland TØ, et al. Triclosan and antimicrobial resistance in bacteria: an overview. Microb Drug Resist. 2006;12: 83–90. doi:10.1089/mdr.2006.12.83

69. Cho I, Yamanishi S, Cox L, Methé B a., Zavadil J, Li K, et al. Antibiotics in early life alter the murine colonic microbiome and adiposity. Nature. 2012;488: 621–626. doi:10.1038/nature11400

70. Loo VG, Bourgault A-M, Poirier L, Lamothe F, Michaud S, Turgeon N, et al. Host and pathogen factors for *Clostridium difficile* infection and colonization. N Engl J Med. 2011;365: 1693–703. doi:10.1056/NEJMoa1012413

71. Thomas C, Stevenson M, Riley T V. Antibiotics and hospital-acquired Clostridium difficile-associated diarrhoea: A systematic review. J Antimicrob Chemother. 2003;51: 1339–1350. doi:10.1093/jac/dkg254

72. Drury B, Scott J, Rosi-Marshall EJ, Kelly JJ. Triclosan exposure increases triclosan resistance and influences taxonomic composition of benthic bacterial communities. Environ Sci Technol. 2013;47: 8923–8930. doi:10.1021/es401919k

73. Meade MJ, Waddell RL, Callahan TM. Soil bacteria Pseudomonas putida and Alcaligenes xylosoxidans subsp. denitrificans inactivate triclosan in liquid and solid substrates. FEMS Microbiol Lett. 2001;204: 45–48. doi:10.1016/S0378-1097(01)00377-9

74. Jones GL, Russell a. D, Caliskan Z, Stickler DJ. A strategy for the control of catheter blockage by crystalline Proteus mirabilis biofilm using the antibacterial agent triclosan. Eur Urol. 2005;48: 838–845. doi:10.1016/j.eururo.2005.07.004

75. Schaffner DW, Bowman JP, English DJ, Fischler GE, Fuls JL, Krowka JF, et al. Quantitative microbial risk assessment of antibacterial hand hygiene products on risk of shigellosis. J Food Prot. 2014;77: 574–82. doi:10.4315/0362-028X.JFP-13-366

76. Birosová L, Mikulásová M. Development of triclosan and antibiotic resistance in Salmonella enterica serovar Typhimurium. J Med Microbiol. 2009;58: 436–441. doi:10.1099/jmm.0.003657-0

77. Ciusa ML, Furi L, Knight D, Decorosi F, Fondi M, Raggi C, et al. A novel resistance mechanism to triclosan that suggests horizontal gene transfer and demonstrates a potential selective pressure for reduced biocide susceptibility in clinical strains of Staphylococcus aureus. Int J Antimicrob Agents. 2012;40: 210–220. doi:10.1016/j.ijantimicag.2012.04.021

78. Mutlu E a, Keshavarzian A, Losurdo J, Swanson G, Siewe B, Forsyth C, et al. A compositional look at the human gastrointestinal microbiome and immune activation parameters in HIV infected subjects. PLoS Pathog. 2014;10: e1003829. doi:10.1371/journal.ppat.1003829

79. Galimanas V, Hall MW, Singh N, Lynch MDJ, Goldberg M, Tenenbaum H, et al. Bacterial community composition of chronic periodontitis and novel oral sampling sites for detecting disease indicators. Microbiome. 2014;2: 32. doi:10.1186/2049-2618-2-32

80. Coyte KZ, Schluter J, Foster KR. The ecology of the microbiome: Networks, competition, and stability. Science. 2015;350: 663–666.

